# Tigmint: Correcting Assembly Errors Using Linked Reads From Large Molecules

**DOI:** 10.1101/304253

**Authors:** Shaun D Jackman ((0000-0002-9275-5966)), Lauren Coombe, Justin Chu, Rene L Warren, Benjamin P Vandervalk, Sarah Yeo, Zhuyi Xue, Hamid Mohamadi, Joerg Bohlmann, Steven JM Jones, Inanc Birol ((0000-0003-0950-7839))

## Abstract

Genome sequencing yields the sequence of many short snippets of DNA (reads) from a genome. Genome assembly attempts to reconstruct the original genome from which these reads were derived. This task is difficult due to gaps and errors in the sequencing data, repetitive sequence in the underlying genome, and heterozygosity, and assembly errors are common. These misassemblies may be identified by comparing the sequencing data to the assembly, and by looking for discrepancies between the two. Once identified, these misassemblies may be corrected, improving the quality of the assembly. Although tools exist to identify and correct misassemblies using Illumina pair-end and mate-pair sequencing, no such tool yet exists that makes use of the long distance information of the large molecules provided by linked reads, such as those offered by the 10x Genomics Chromium platform. We have developed the tool Tigmint for this purpose. To demonstrate the effectiveness of Tigmint, we corrected assemblies of a human genome using short reads assembled with ABySS 2.0 and other assemblers. Tigmint reduced the number of misassemblies identified by QUAST in the ABySS assembly by 216 (27%). While scaffolding with ARCS alone more than doubled the scaffold NGA50 of the assembly from 3 to 8 Mbp, the combination of Tigmint and ARCS improved the scaffold NGA50 of the assembly over five-fold to 16.4 Mbp. This notable improvement in contiguity highlights the utility of assembly correction in refining assemblies. We demonstrate its usefulness in correcting the assemblies of multiple tools, as well as in using Chromium reads to correct and scaffold assemblies of long single-molecule sequencing. The source code of Tigmint is available for download from https://github.com/bcgsc/tigmint, and is distributed under the GNU GPL v3.0 license.

## Introduction

Assemblies of short read sequencing data are easily confounded by repetitive sequences larger than the fragment size of the sequencing library. When the size of a repeat exceeds the library fragment size, the contig comes to an end in the best case, or results in misassembled sequence in the worst case. Misassemblies not only complicate downstream analyses, but also limit the contiguity of the assembly, when incorrectly assembled sequences prevent joining their adjacent and correctly assembled sequences during assembly scaffolding.

Long-read sequencing technologies have greatly improved assembly contiguity, by their ability to span these repeats, but at a cost roughly ten times that of short-read sequencing technology. For population studies and when sequencing large genomes, such as conifer genomes and other economically important crop species, this cost may be prohibitive. The 10x Genomics (Pleasanton, CA) Chromium technology generates linked reads from large DNA molecules at a cost comparable to standard short-read sequencing technologies. Whereas paired-end sequencing gives two reads from a small DNA fragment, linked reads yield roughly a hundred read pairs from molecules with a typical size of a hundred kilobases. Linked reads indicate which reads were derived from the same DNA molecule (or molecules, when they share the same barcode), and so should be in close proximity in the underlying genome. The technology has been used previously to phase diploid genomes using a reference [1], *de novo* assemble complex genomes in the gigabase scale [2], and further scaffold draft assemblies [3].

A number of software tools employ linked reads for various applications. The LongRanger tool maps reads to repetitive sequence, phase small variants, and identify structural variants (https://www.10xgenomics.com/software/), and Supernova [2] assembles diploid genome sequences, both tools developed by the vendor. Among tools from academic labs, GROC-SVs [4], NAIBR [5], and Topsorter (https://github.com/hanfang/Topsorter) identify structural variants, and ARCS [6], Architect [7], and fragScaff [8] scaffold genome assemblies using linked reads.

In *de novo* sequencing projects, it is challenging yet important to ensure the correctness of the resulting assemblies. Tools to correct misassemblies typically inspect the reads aligned back to the assembly to identify discrepancies. Pilon [9] inspects the alignments to identify variants and correct small-scale misassemblies. NxRepair [10] uses Illumina mate-pair sequencing to correct large-scale structural misassemblies. Linked reads offer an opportunity to use the long-range information provided by large molecules to identify misassemblies, yet no software tool currently exists to correct misassemblies using linked reads. Here we introduce a software tool, Tigmint, to identify misassemblies using this new and useful data type.

Tigmint first aligns reads to the assembly, and infers the extents of the large DNA molecules from these alignments. It then searches for atypical drops in physical molecule coverage. Since the physical coverage of the large molecules is more consistent and less prone to coverage dropouts than that of the short read sequencing data, it can be used to reveal the positions of possible misassemblies. Linked reads may be used to scaffold the corrected assembly with ARCS [6] to identify contig ends sharing barcodes, and either ABySS-Scaffold (included with ABySS) or LINKS [11] to merge sequences of contigs into scaffolds.

## Methods

### Algorithm

The user provides a draft assembly in FASTA format and the reads in FASTQ format. Tigmint first aligns the reads to the draft genome using BWA-MEM [12]. The alignments are filtered by alignment score and number of mismatches to remove poorly aligned reads with the default thresholds NM < 5 and AS ≥ 0.65 · *l*, where *l* is the read length. Reads with the same barcode that map within 50 kbp of the adjacent read are grouped into a molecule and assigned a unique numeric molecule identifier. A tab-separated values (TSV) file is constructed, where each record indicates the start and end of one molecule, and records the number of reads that compose that molecule, their median mapping quality, alignment score, and number of mismatches. Unusually small molecules, shorter than 2000 bp by default, are filtered out.

Physical molecule depth of coverage counts the number of molecules that span a point. Regions with poor physical molecule coverage indicate potentially problematic regions of the assembly. At a misassembly involving a repeat, molecules may start in left flanking unique sequence and end in the repeat, and molecules may start in the repeat and end in right flanking unique sequence. This seemingly uninterrupted molecule coverage may give the appearance that the region is well covered by molecules. Closer inspection may reveal that no molecules span the repeat entirely, from the left flanking sequence to the right flanking sequence. Tigmint checks that each region of a fixed size specified by the user, 2000 bp by default, is spanned by a minimum number of molecules, 20 by default. The Python package Intervaltree is used to efficiently identify regions with insufficient spanning molecules. Regions with fewer spanning molecules reveal possible misassemblies, and the locations of these regions are written to a BED file. The sequences of the original draft assembly are cut at these breakpoints, producing a corrected FASTA file.

Tigmint will optionally run ARCS [6] to scaffold these corrected sequences and improve the contiguity of the assembly. Tigmint will optionally compare the scaffolds to a reference genome, if one is provided, using QUAST [13] to compute contiguity (NGA50) and correctness (number of misassemblies) of the assemblies before Tigmint, after Tigmint, and after ARCS. Each misassembly identified by QUAST indicates a difference between the assembly and the reference. These putative misassemblies are composed of both misassemblies and structural variation between the reference and the sequenced genomes. The NGA50 metric summarizes both assembly contiguity and correctness by computing the NG50 of the lengths of alignment blocks to a reference genome, correcting the contiguity metric by accounting for possible misassemblies. It however also penalizes sequences at points of true variation between the sequenced and reference genomes. The true but unknown contiguity of the assembly, which accounts for misassemblies but not for structural variation, therefore lies somewhere between the lower bound of NGA50 and the upper bound of NG50.

### Correcting human assemblies

We downloaded the ABySS 2.0 [14] assembly `abyss-2.0/scaffolds.fa` from http://bit.ly/giab-hg004 for the Genome in a Bottle (GIAB) HG004, assembled from Illumina paired-end and mate-pair reads [15]. We downloaded the 10x Genomics Chromium reads for this same individual from http://bit.ly/giab-hg004-chromium and used the LongRanger Basic pipeline to extract the barcodes from the reads. We ran Tigmint to correct the ABySS 2.0 assembly of HG004 using these Chromium reads with the parameters window = 2000 and span = 20. The choice of parameters is discussed in the results. Both the uncorrected and corrected assemblies are scaffolded using ARCS. These assemblies are compared to the chromosome sequences of the GRCh38 reference genome using QUAST [13]. Since ARCS does not estimate gap sizes using linked reads, the QUAST parameter `—scaffold-gap-max-size` is set to 100 kbp.

We repeated this analysis using Tigmint, ARCS, and QUAST, with five other assemblies. We downloaded from http://bit.ly/giab-hg004the reads assembled with DISCOVARdenovo and scaffolded using BESST [16], and the same DISCOVARdenovo contigs scaffolded using ABySS-Scaffold. We assembled the linked reads with Supernova 2.0.0 [2], which used neither the 2×250 paired-end reads nor mate-pair reads. We applied Tigmint and ARCS to two assemblies of single-molecule sequencing (SMS) reads. We downloaded PacBio reads assembled with Falcon from http://bit.ly/giab-falcon [17] and Oxford Nanopore reads assembled with Canu [18], accession GCA_900232925.1. The Canu assembly is of NA12878, and all other assemblies are of NA24143 (HG004). The script to run this analysis is available online at https://github.com/sjackman/tigmint-data. Most software used in these analyses were installed using Linuxbrew [19] with the command `brew install abyss arcs bwa lrsim miller minimap2 samtools seqtk`. We used the development version of QUAST 5 revision 78806b2, which is capable of analyzing assemblies of large genomes using Minimap2 [20].

## Results

Correcting the ABySS assembly of the human data set HG004 with Tigmint reduces the number of misassemblies identified by QUAST by 216, a reduction of 27%. While the scaffold NG50 decreases slightly from 3.65 Mbp to 3.47 Mbp, the scaffold NGA50 remains unchanged; thus in this case, correcting the assembly with Tigmint improves the correctness of the assembly without substantially reducing its contiguity. However, scaffolding the uncorrected and corrected assemblies with ARCS yield markedly different results: a 2.5-fold increase in NGA50 from 3.1 Mbp to 7.9 Mbp without Tigmint versus a more than five-fold increase in NGA50 to 16.4 Mbp with Tigmint. Further, correcting the assembly and then scaffolding yields a final assembly that is both more correct and more contiguous than the original assembly, as shown in Fig. 1 and Table 1.

**Figure 1:**
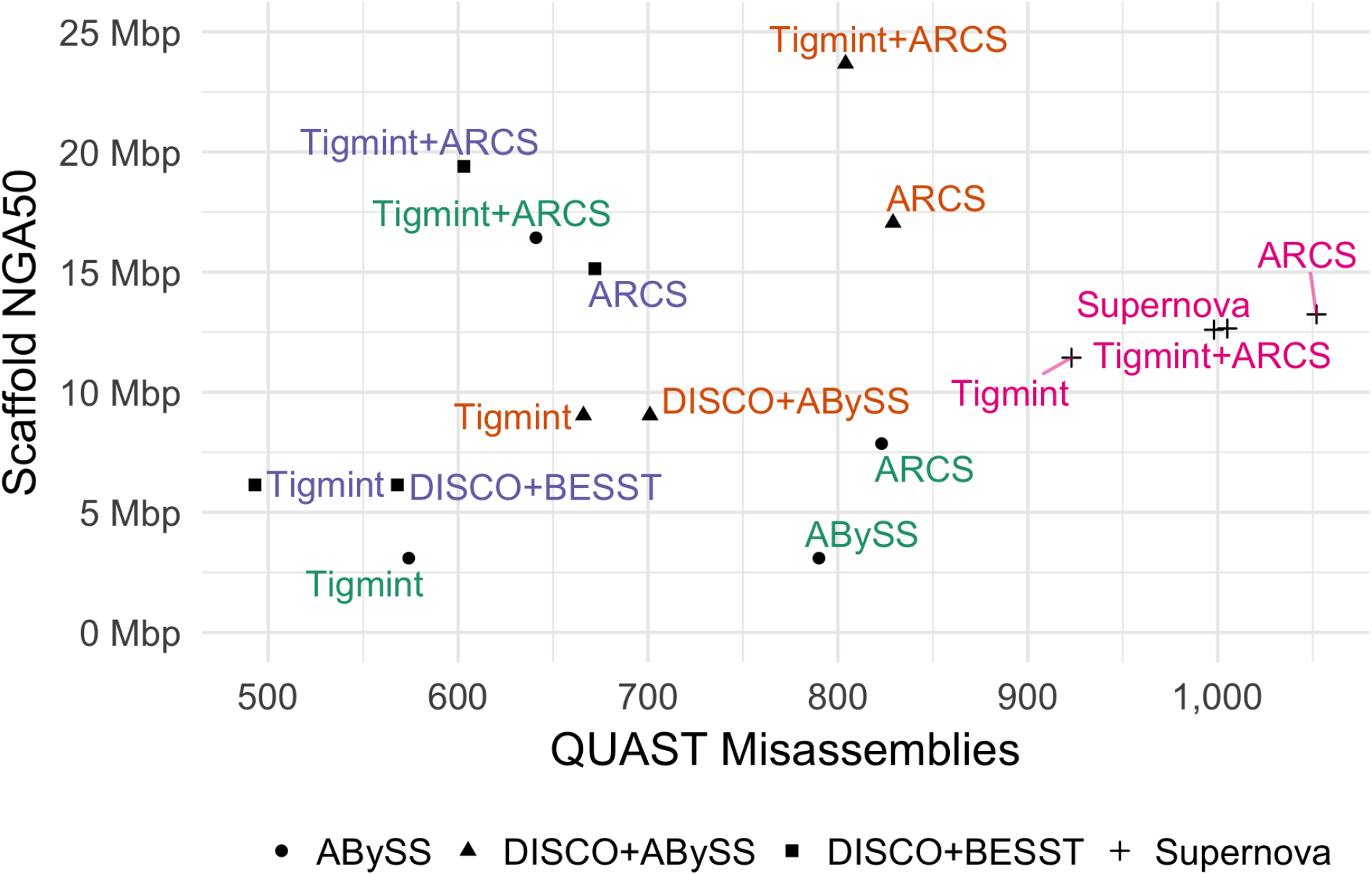
Assembly contiguity and correctness metrics with and without correction using Tigmint prior to scaffolding with ARCS. The most contiguous and correct assemblies are found in the top-left. Supernova used linked reads only, whereas the others used paired end and mate pair reads.

**Table 1:** The assembly contiguity (scaffold NG50 and NGA50) and correctness (number of misassemblies) metrics with and without correction using Tigmint prior to scaffolding with ARCS.

| Assembly | NG50 (Mbp) | NGA50 (Mbp) | Misassemblies | Reduction |
| --- | --- | --- | --- | --- |
| ABySS | 3.65 | 3.09 | 790 | NA |
| ABySS+Tigmint | 3.47 | 3.09 | 574 | 216 (27.3%) |
| ABySS+ARCS | 9.91 | 7.86 | 823 | NA |
| ABySS+Tigmint+ARCS | 26.39 | 16.43 | 641 | 182 (22.1%) |
| DISCO+ABySS | 10.55 | 9.04 | 701 | NA |
| DISCO+ABySS+Tigmint | 10.16 | 9.04 | 666 | 35 (5.0%) |
| DISCO+ABySS+ARCS | 29.20 | 17.05 | 829 | NA |
| DISCO+ABySS+Tigmint+ARCS | 35.31 | 23.68 | 804 | 25 (3.0%) |
| DISCO+BESST | 7.01 | 6.14 | 568 | NA |
| DISCO+BESST+Tigmint | 6.77 | 6.14 | 493 | 75 (13.2%) |
| DISCO+BESST+ARCS | 27.64 | 15.14 | 672 | NA |
| DISCO+BESST+Tigmint+ARCS | 33.43 | 19.40 | 603 | 69 (10.3%) |
| Supernova | 38.48 | 12.65 | 1,005 | NA |
| Supernova+Tigmint | 17.72 | 11.43 | 923 | 82 (8.2%) |
| Supernova+ARCS | 39.63 | 13.24 | 1,052 | NA |
| Supernova+Tigmint+ARCS | 27.35 | 12.60 | 998 | 54 (5.1%) |
| Falcon | 4.56 | 4.21 | 3,640 | NA |
| Falcon+Tigmint | 4.45 | 4.21 | 3,444 | 196 (5.4%) |
| Falcon+ARCS | 18.14 | 9.71 | 3,801 | NA |
| Falcon+Tigmint+ARCS | 22.52 | 11.97 | 3,574 | 227 (6.0%) |
| Canu | 7.06 | 5.40 | 1,688 | NA |
| Canu+Tigmint | 6.87 | 5.38 | 1,600 | 88 (5.2%) |
| Canu+ARCS | 19.70 | 10.12 | 1,736 | NA |
| Canu+Tigmint+ARCS | 22.01 | 10.85 | 1,626 | 110 (6.3%) |
| Simulated | 9.00 | 8.28 | 272 | NA |
| Simulated+Tigmint | 8.61 | 8.28 | 217 | 55 (20.2%) |
| Simulated+ARCS | 23.37 | 17.09 | 365 | NA |
| Simulated+Tigmint+ARCS | 30.24 | 24.98 | 320 | 45 (12.3%) |

Correcting the DISCOVARdenovo + BESST assembly reduces the number of misassemblies by 75, a reduction of 13%. Using Tigmint to correct the assembly before scaffolding with ARCS yields an increase in NGA50 of 28% over using ARCS without Tigmint. Correcting the DISCOVARdenovo + ABySS-Scaffold assembly reduces the number of misassemblies by 35 (5%), after which scaffolding with ARCS improves the NGA50 to 23.7 Mbp, 2.6 times the original assembly and a 40% improvement over ARCS without Tigmint. Three assemblies are on the Pareto frontier maximizing NGA50 and minimizing misassemblies. The assembly with the fewest misassemblies is DISCOVAR + BESST + Tigmint. The assembly with the largest NGA50 is DISCOVAR + ABySS-Scaffold + Tigmint + ARCS. The last assembly on the Pareto frontier is DISCOVARdenovo + BESST + Tigmint + ARCS, which strikes a good balance between both good contiguity and few misassemblies.

Correcting the Supernova assembly of the HG004 linked reads with Tigmint reduces the number of misassemblies by 82, a reduction of 8%, and after scaffolding the corrected assembly with ARCS, we see a slight (<1%) decrease in both misassemblies and NGA50 compared to the original Supernova assembly. Since the Supernova assembly is composed entirely of the linked reads, we do not expect significant gains from using these same data to correct and scaffold the Supernova assembly. The Supernova assembly however has not made use of the mate-pair reads, and correcting the Supernova assembly with mate-pair reads may be an interesting area for future development of Tigmint.

The assemblies of SMS reads have contig NGA50s in the megabases. Tigmint and ARCS together improve the scaffold NGA50 of the Canu assembly by more than double to nearly 11 Mbp and improve the scaffold NGA50 of the Falcon assembly by nearly triple to 12 Mbp, and both assemblies have fewer misassemblies than their original assembly, shown in Fig. 2. Using Tigmint and ARCS together improves both the contiguity and correctness over the original assembly. Using linked reads in combination with long reads, we can achieve an assembly that achieves both a high contig NGA50 as well as high scaffold NGA50, which is not currently possible with either technology alone.

**Figure 2:**
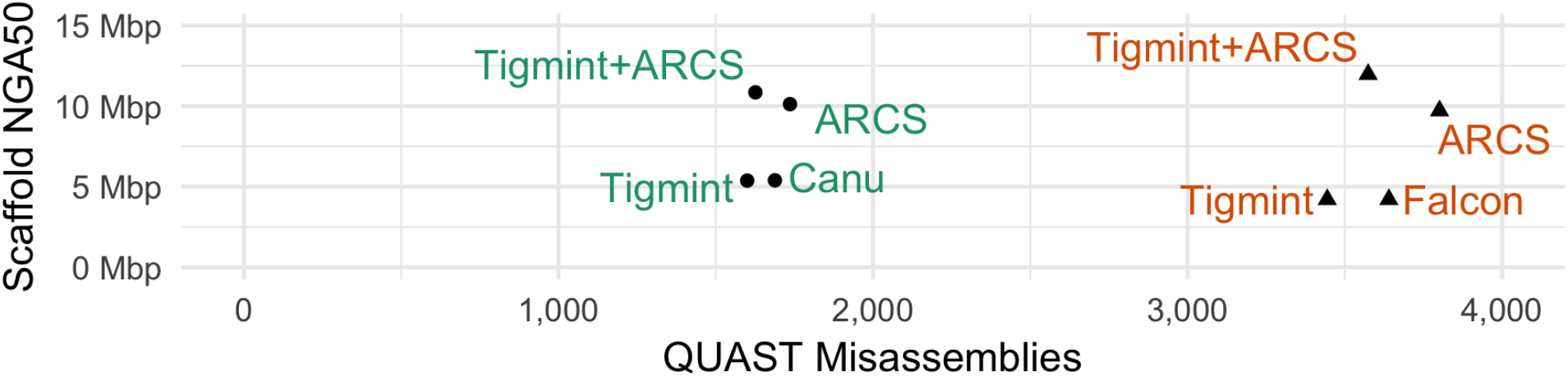
Assemblies of Nanopore reads with Canu and PacBio reads with Falcon with and without correction using Tigmint prior to scaffolding with ARCS.

The alignments of the ABySS assembly to the reference genome before and after Tigmint are visualized in Fig. 3 using JupiterPlot (https://github.com/JustinChu/JupiterPlot). A number of split alignments, likely misassemblies, are visible in the assembly before Tigmint, whereas after Tigmint no such split alignments are visible.

**Figure 3:**
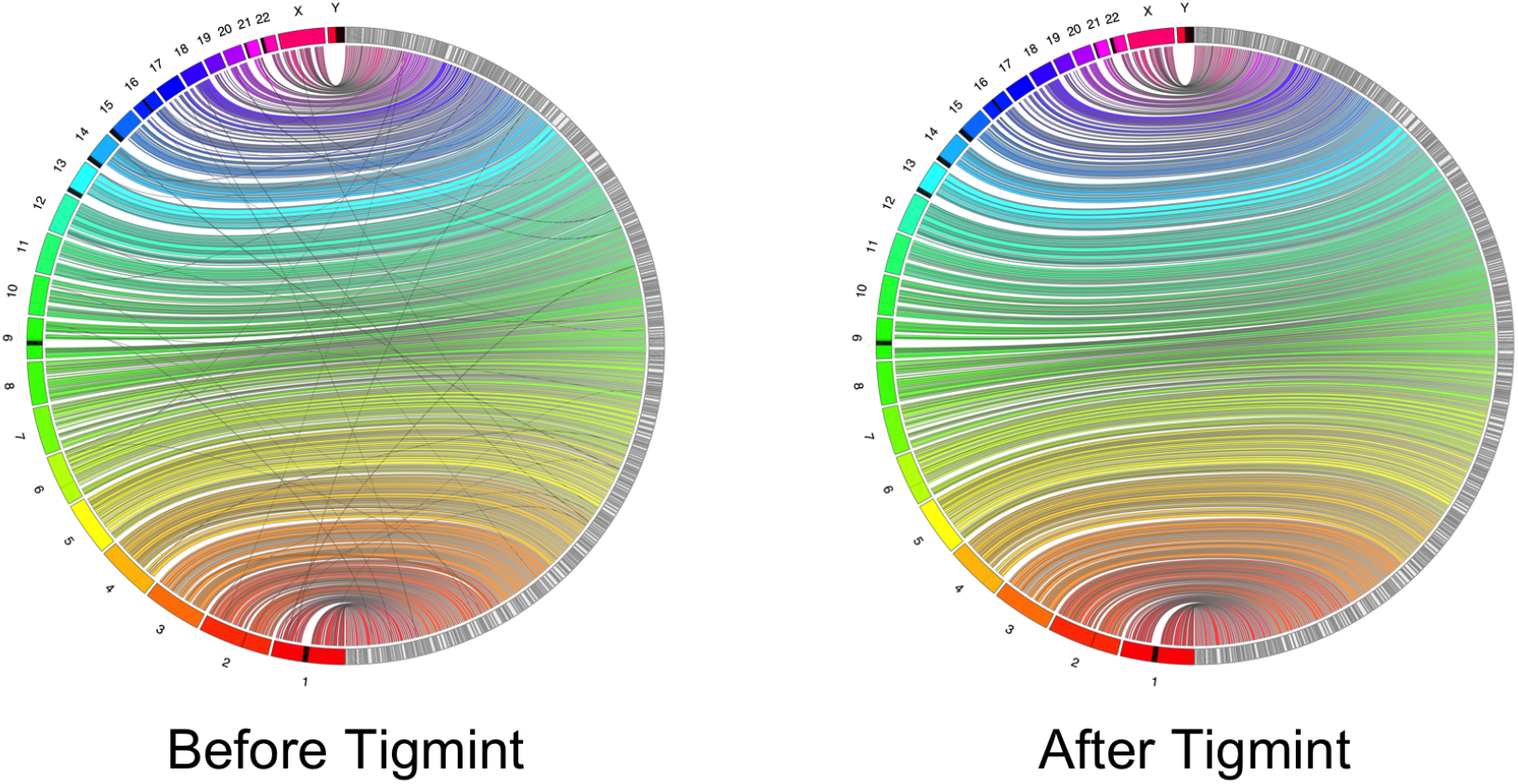
The alignments to the reference genome of the ABySS assembly before and after Tigmint. The reference chromosomes are on the left in colour, the assembly scaffolds on the right in grey. No translocations are visible after Tigmint.

The default maximum distance permitted between linked reads in a molecule is 50 kbp, which is the value used by the Longranger and Lariat tools of 10x Genomics. Values between 20 kbp and 100 kbp do not substantially affect the results, and values smaller than 20 kbp begin to disconnect linked reads which should be found in a single molecule. The effect of varying the window and spanning molecules parameters of Tigmint on the assembly contiguity and correctness metrics is shown in Fig. 4. The assembly metrics of the ABySS, DISCOVARdenovo + ABySS-Scaffold, and DISCOVARdenovo + BESST assemblies after correction with Tigmint are rather insensitive to the spanning molecules parameter for any value up to 50 and for the window parameter for any value up to 2 kbp. The DISCOVAR + BESST assembly had the fewest misassemblies when span ≥ 20 and window ≥ 2000. Larger values for either of these parameters yielded for the ABySS and DISCOVARdenovo + ABySS assemblies either more misassemblies or lower NGA50. Based on these results, we selected default values of span = 20 and window = 2000, which worked well for all of the tested assembly tools. When varying the spanning molecules parameter, the window parameter is fixed at 2000, and when varying the window parameter, the spanning molecules parameter is fixed at 20. The median molecule depth is 163, computed using Bedtools [21], and its inter-quartile range (IQR) is 31.

**Figure 4:**
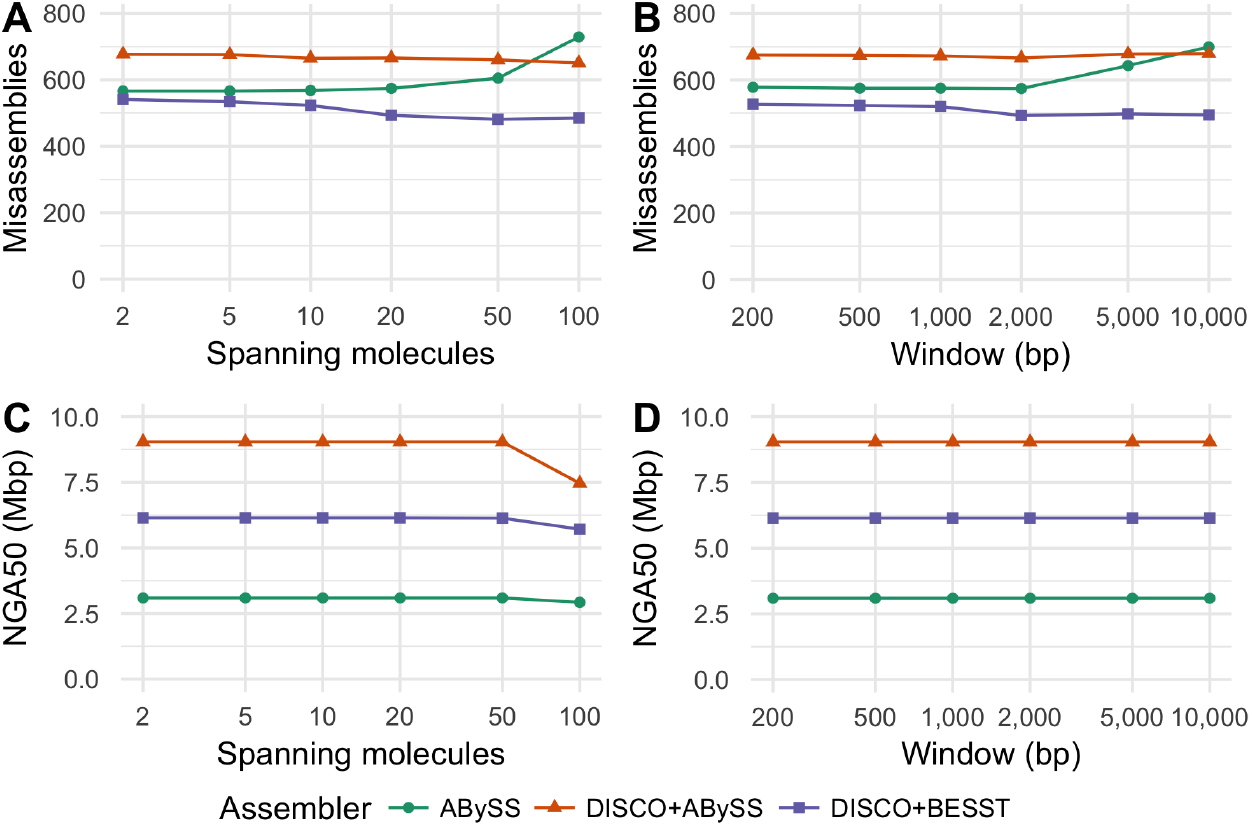
Effect of varying the window and span parameters on scaffold NGA50 and misassemblies.

We simulated 434 million 2×250 paired-end and 350 million 2×125 mate-pair read pairs using wgsim of samtools and 524 million 2×150 linked read pairs using LRSim [22], emulating the HG004 data set. We assembled these reads using ABySS 2.0.2, and applied Tigmint and ARCS as before. The assembly metrics are shown in Table 1. We see similar performance to the real data, a 20% reduction in misassemblies after running Tigmint, and a three-fold increase in NGA50 after Tigmint and ARCS. Since no structural rearrangements are present in the simulated data, each misassembly identified by QUAST ought to be a true misassembly, allowing us to calculate precision and recall. For the parameters used with the real data, window = 2000 and span = 20, Tigmint makes 210 cuts in scaffolds at least 3 kbp (QUAST does not analyze shorter scaffolds), and corrects 55 misassemblies of the 272 identified by QUAST, yielding precision and recall of PPV = 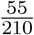 = 0.26 and TPR = 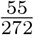 = 0.20. Using instead window = 1000, Tigmint makes only 58 cuts and yet corrects 51 misassemblies, making its precision and recall PPV = 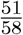 = 0.88 and TPR = 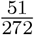 = 0.19, a marked improvement in precision with only a small decrease in recall. The scaffold NGA50 after ARCS is 24.7 Mbp, 1% less than with window = 2000. Since the final assembly metrics are similar, using a smaller value for the window size parameter may avoid unnecessary cuts. Small-scale misassemblies cannot be detected by Tigmint, such as collapsed repeats, and relocations and inversions smaller than a typical molecule. We intend to investigate the remaining misassemblies identified by QUAST to assess their nature, and determine whether some could be detected by Tigmint with further development.

The primary steps of running Tigmint are mapping the reads to the assembly, determining the start and end coordinate of each molecule, and finally identifying the discrepant regions and correcting the assembly. Mapping the reads to the DISCOVAR + ABySS-Scaffold assembly with BWA-MEM and concurrently sorting by barcode using Samtools [23] in a pipe required 5.5 hours (wall-clock) and 17.2 GB of RAM (RSS) using 48 threads on a 24-core hyper-threaded computer. Determining the start and end coordinates of each molecule required 3.25 hours and 0.08 GB RAM using a single thread. Finally, identifying the discrepant regions of the assembly, correcting the assembly, and creating a new FASTA file required 7 minutes and 3.3 GB RAM using 48 threads. The slowest step of mapping the reads to the assembly could be made faster by using light-weight mapping rather than full alignment, since Tigmint needs only the positions of the reads, not their alignments. The speed of determining the start and end coordinates of each molecule could likely be improved with either multithreading or by rewriting this tool, currently implemented in Python, in C++. The final step of identifying the discrepant regions of the assembly is implemented in Python and is not a bottle neck.

## Discussion

When aligning an assembly of an individual’s genome to a reference genome of its species, we expect to see breakpoints where the assembled genome differs from the reference genome. These breakpoints are caused by both misassemblies and true differences between the individual and the reference. The median number of mobile-element insertions for example, just one class of structural variant, is estimated to be 1,218 per individual [24]. Misassemblies can be corrected by inspecting the alignments of the reads to the assembly and cutting the scaffolds at positions not supported by the reads. Misassemblies due to true structural variation will however remain. For this reason, even a perfectly corrected assembly is expected to have a number of differences when compared to the reference.

Tigmint uses linked reads to reduce the number of misassemblies in a genome sequence assembly. The contiguity of the assembly is not appreciably affected by such a correction, while yielding an assembly that is more correct. Most scaffolding tools order and orient the sequences that they are given, but do not attempt to correct misassemblies. These misassemblies hold back the contiguity that can be achieved by scaffolding. Two sequences that should be connected together cannot be when one of those two sequences is connected incorrectly to a third sequence. By first correcting these misassemblies, the scaffolding tool can do a better job of connecting sequences, and we observe precisely this harmonious effect. Scaffolding an assembly that has been corrected with Tigmint yields a final assembly that is both more correct and substantially more contiguous than an assembly that has not been corrected.

Using single-molecule sequencing in combination with linked reads enables a genome sequence assembly that achieves both a high sequence contiguity as well as high scaffold contiguity, a feat not currently achievable with either technology alone. Although high-throughput short-read sequencing is often used to polish a long-read assembly to improve its accuracy at the nucleotide level, short read sequencing reads align ambiguously to repetitive sequence, and so are not well suited to polish the repetitive sequence of the assembly. Linked reads would resolve this mapping ambiguity and are uniquely suited to polishing an assembly of long reads, an opportunity for further research in the hybrid assembly of long and linked reads.

